# A *Candidozyma* (*Candida*) *auris*-optimized Episomal Plasmid Induced Cas9-editing system reveals the direct impact of the S639F encoding *FKS1* mutation

**DOI:** 10.1101/2025.02.14.638356

**Authors:** Laura A. Doorley, Vanessa Meza-Perez, Sarah J. Jones, Jeffrey M. Rybak

**Affiliations:** Department of Pharmacy and Pharmaceutical Sciences, St. Jude Children’s Research

**Keywords:** *Candida auris*, CRISPR, EPIC, gene editing, resistance, *FKS1*

## Abstract

**Objectives:** Mutations in the *Candidozyma* (*Candida*) *auris* β-glucan synthase gene (*FKS1*) altering S639 are frequently associated with clinical echinocandin resistance. However, the direct impact of these mutations remains uncharacterized. We have developed a novel *C. auris-*optimized **E**pisomal **P**lasmid-**I**nduced **C**as9 (EPIC) gene-editing system capable of recyclable precision genome editing and demonstrate the contribution of *FKS1*^S639F^ mutation to echinocandin resistance.

**Methods:** The EPIC gene-editing system was generated for optimized use in *C. auris*, and *ADE2* modification was evaluated in 5 *C. auris* clades. Mutations leading to Fks1^S639F^ and Fks1^WT^ were placed into echinocandin-susceptible and echinocandin-resistant isolates from Clade-III and -I, respectively, using the EPIC system. Echinocandin susceptibility was determined by CLSI broth microdilution, and cell wall abundance of chitin and β-glucan was assessed by staining with Calcofluor White (CFW) and Aniline Blue (AB).

**Results:** The EPIC system was capable of targeted *ADE2* editing in *C. auris* isolates from 5 genetic clades and shown to be precise by confirmatory sequencing of 50 transformants. A single nucleotide change in *FKS1* resulting in either the S639F substitution or a silent synonymous mutation was introduced in an echinocandin-susceptible Clade-III isolate.

Precision *FKS1* editing by the EPIC system was confirmed by whole genome sequencing. Subsequent susceptibility testing demonstrated introduction of the *FKS1* S639F mutation to increase resistance to echinocandins. Moreover, introduction of a wildtype Fks1 sequence to an echinocandin-resistant Clade-I clinical isolate, correcting the *FKS1*^S639F^ mutation, resulted in a restoration of echinocandin sensitivity. Evaluation of cell wall composition showed isolates or strains harboring *FKS1*^S639F^ to contain significantly elevated β-glucan and chitin content relative to isogenic comparators.

**Conclusions:** These data demonstrate the potential of our EPIC system in its ability to introduce single nucleotide substitutions in multiple *C. auris* clade backgrounds while revealing the direct impact of the S639F encoding *FKS1* mutation on echinocandin resistance.

## INTRODUCTION

*C. auris* is an emerging fungal pathogen recognized by the Centers for Disease Control and Prevention (CDC) as an increasing threat to public health, calling for “urgent and aggressive action” [1, 2]. Applying the current tentative CDC clinical breakpoints, approximately 90% of *C. auris* clinical isolates are found to be resistant to fluconazole, and up to 50% of isolates are resistant to amphotericin B [3]. Typically, echinocandin resistance is observed in only 1 to 5% of isolates, making echinocandins the front-line therapy of choice for many clinicians [3]. However, numerous cases of *C. auris* infections acquiring echinocandin resistance in host during treatment have been documented, leaving clinicians with no reliable therapeutic option when treating patients with *C. auris* infections [4, 5].

Previous studies have shown that echinocandin resistance in species, such as *Candida glabrata* and *Candida albicans* is strongly associated with mutations in the genes encoding the echinocandin target β-glucan synthase (*GSC1*, also known as *FKS1*, in *C. albicans* and the genes *FKS1* and *FKS2* in *C. glabrata*) [6-8]. Furthermore, these mutations can be predictive of clinical failure for candidemia treatment [9-11]. The vast majority of *FKS1* mutations cluster into 3 distinct “hot-spots”, all of which encode residues spatially near a single hydrophobic convex surface predicted to interact with both plasma membrane lipids and echinocandins [7, 12-14] While the association between these mutations and clinical echinocandin resistance is undeniable, confirmation of the direct impact of these β-glucan synthase mutations by genetic interrogation has been hindered both by limited genetic tools and the practicality of manipulating the gene of interest (essentiality, large size: approximately 5.7kb, and proximity to flanking features). Previous characterization of β-glucan synthase mutations has relied on methods requiring exposure of cells to echinocandins (possibly introducing confounding adaptations) either directly through *in vitro* evolution studies or indirectly as a component of selection media to identify potential transformants [15, 16] Alternatively, clinically derived *Candida FKS1* mutations have been recreated *in situ* with the addition of large selection cassettes or through heterologous expression studies by site-directed mutagenesis in the model organism *Saccharomyces cerevisiae* [7, 17]. Consequently, studies demonstrating the direct impact of mutations in *Candida* β-glucan synthase on clinical echinocandin resistance without these potential confounding variables are limited.

In this work, we describe the creation of the *C. auris-*optimized **E**pisomal **P**lasmid **I**nduced **C**as9 (EPIC) genetic manipulation system, a transient and recyclable tool capable of single nucleotide editing without the need to integrate a dominant marker into the *C. auris* genome. We demonstrate system functionality in clinical isolates representing different clades of *C. auris*, confirm system precision using whole genome sequencing (WGS), and utilize the system to interrogate direct impacts of the *FKS1* mutation leading to S639F substitution in two distinct *C. auris* clinical isolate backgrounds.

## MATERIALS AND METHODS

### Isolate, strains, and growth media

All clinical isolates and derived strains used in this study are listed in **Supplementary Table 1**. Long-term *C. auris* isolates and strains were maintained in 40% glycerol at -80°C and cultured in YPD media (1% yeast extract, 2% peptone, 2% dextrose) at 35°C, 220 RPM, unless noted otherwise.

### Construction of EPIC system components

The *C. auris-*optimized EPIC vector, pJMR19 (accession: *pending*) was constructed from pJMR17v3 utilizing restriction digest and ligation to remove the *C. albicans* ORI410 sequence and to replace the *C. albicans ACT1* promoter, with the 1kb sequence upstream of the *C. auris ACT1* gene (B9J08_000486) [18]. Vector verification via whole plasmid sequencing was performed by Plasmidsaurus using Oxford Nanopore Technology with custom analysis and annotation.

Duplexed guide sequence primers targeting the gene of interest were ligated into pJMR19 digested by Lgu1 (Thermo Scientific™), as previously described [18, 19]. Transformation repair templates were created by PCR amplification of gBlock sequence (Integrated DNA technologies) followed by QIAquick PCR Purification Kit (Qiagen) or by duplexing approximately 80bp complementary primers (Integrated DNA Technologies) as previously described [18, 19]. All vectors, primers, and templates are listed in **Supplementary Table 1**.

### EPIC-based *C. auris* transformation

*C. auris* was grown overnight in 25mL YPD to an OD_600_ of 1.8-2.2. Cultures were pelleted at 4000 RPM for three minutes, washed with 1mL water, and pellets were resuspended in 1mL TE-LiAc (1M Tris-EDTA and 1M lithium acetate pH7.5) and centrifuged once more at 8000 RPM for 30 seconds. Pellets were resuspended in a minimally required TE-LiAc volume.

Transformation reactions were assembled with 10µL boiled then cooled salmon sperm DNA, 8µg pJMR19, 8µg repair template DNA, 50µL resuspended pellet, and 300µL TE-LiAC + 55% polyethylene glycol (PEG). Transformations were incubated for 90 minutes with intermittent manual agitation, heat-shocked at 45°C for 15 minutes and pelleted at 8000 RPM for 30 seconds. Pellets were resuspended in 2mL YPD and incubated for 4 hours. Transformants were cultured on nourseothricin (200µg/mL)-supplemented YPD agar plates for selection. Single colonies were patched, screened and replica plated for plasmid ejection as previously described [18, 19].

### Validation of EPIC manipulation and specificity

Genomic DNA (gDNA) was extracted from overnight *C. auris* transformant cultures using the MasterPure™ Yeast DNA Purification kit (Biosearch Technologies). *C. auris FKS1* (B9J08_000964) and ‘landing-pad’ sequences were amplified by PCR and purified with QIAquick PCR Purification Kits (Qiagen). *FKS1* sequencing was performed by Plasmidsaurus using Oxford Nanopore Technology. *C. auris ADE2* (B9J08_03951) deep sequencing was performed by the Center for Advanced Genome Engineering at St. Jude Children’s Research Hospital (SJCRH) utilizing a two-step PCR system and the Illumina Miseq sequencing platform.

WGS was performed for the clinical isolate RVA001 and select RVA001-derived strains. Sample libraries were prepared from gDNA by the Hartwell Center at SJCRH using KAPA HyperPrep (Roche) and sequencing targeting 100x coverage/ 150bp reads was performed via Illumina HiSeq. Code4DNA (Code4DNA.com) provided bioinformatics analysis. Two sequence library runs per sample were merged for each paired end R1 and R2. Overlapping regions were error corrected using bbmerge 38.94. Reads were aligned to GCA_002775015.1 using bwa (0.7.17-r1188). Alignments were sorted and indexed by Samtools v1.13 and duplicates were marked with picard (2.26.0). Variants called by running chromosomal based Freebayes (1.3.4) on GNU parallel (20180722) were annotated using SnpEff (4.5covid19). Variants found in each affected sample but not the control were extracted with Vcftools (0.1.15) and manually evaluated using Jbrowse (jbrowse-desktop-v1.5.3). (SRA: *pending*).

### Antifungal susceptibility testing

Minimum inhibitory concentrations (MIC) for anidulafungin (Sigma-Aldrich), caspofungin (Fisher Scientific), micafungin (Sigma-Aldrich), and rezafungin (MedChemExpress) were determined by broth microdilution in accordance with the M27-A4 from the Clinical Laboratory Standards Institute [20]. All susceptibility testing was performed in biological triplicate.

### Relative cell wall constituent abundance

*C. auris* strains grown on Sabouraud-dextrose (SD) agar were subsequently cultured in RPMI (RPMI1640, MOPS, 2% glucose, pH7) to mid-log phase. Suspensions were washed to remove media and fixed with 3.7% (v/v) formaldehyde. Pellets were washed, normalized for cell density, aliquoted, and stained for chitin or β-glucans with 25mg/L CFW (Fluorescent Brightener 28: Fisher Scientific) or 50mg/L AB (Fisher Scientific), respectively. BioTek Cytation7 with Gen5 software (Agilent) used to measure absorbance (OD_600_) and fluorescence intensity (CFW: 380nm/475nm; AB: 395nm/495nm).

## RESULTS

### The EPIC genetic manipulation system is functional in multiple *C. auris* clinical isolate backgrounds

The *C. auris* gene B9J08_03951 (*ADE2*), predicted to encode phosphoribosylaminoimidazole carboxylase, was selected to demonstrate the functionality of the EPIC genetic manipulation system. Successful *ADE2* disruption (*ADE2*^dis^) produces red-pink colony coloration on standard culture media (SD or YPD) due to accumulation of pigmented metabolites in the adenine biosynthetic pathway **(Figure 1)**. *ADE2*^dis^ transformations were carried out with the EPIC vector containing a guide sequence targeting the 5’ region of *ADE2* (pJMR19-*dADE2*) and a DNA repair template encoding approximately 1000 bases of the 5’ region of *ADE2* in five clinical isolates representing five *C. auris* clades. Transformants from each background including RVA001 (Clade-III, *FKS1*^WT^) and SKU067 (Clade-I, *FKS1*^S639F^), the *C. auris* isolates to be studied in subsequent *FKS1-*related experiments, yielded red-pink colonies. (**Supplementary Figure 1**).

**Figure 1.**
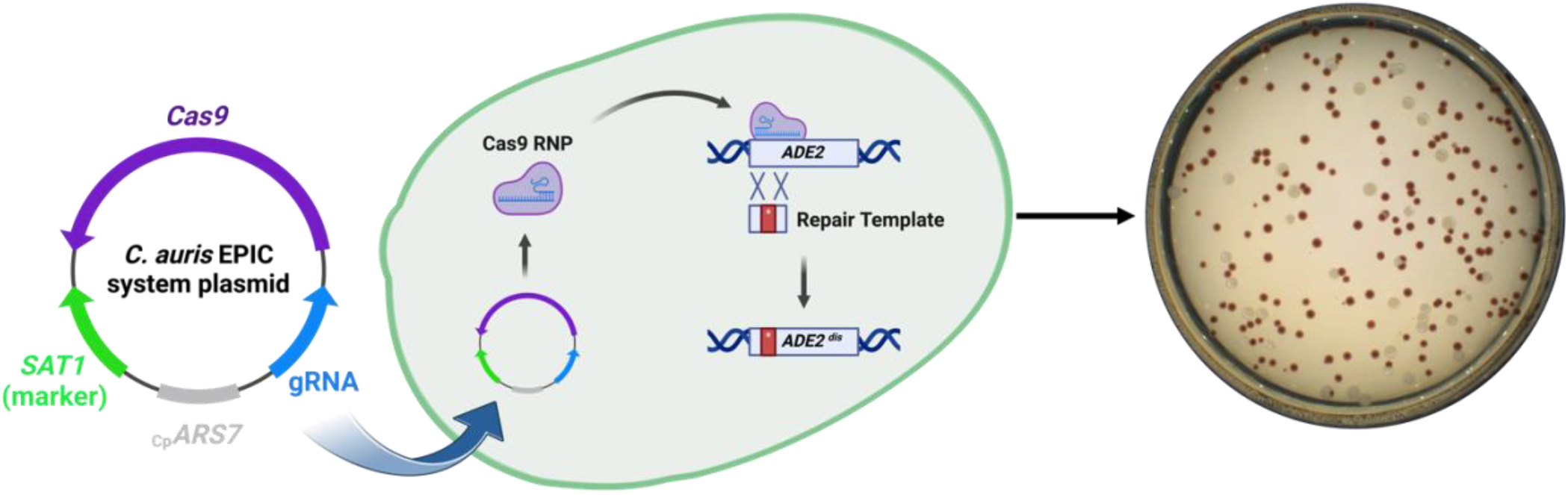
Genetic manipulation of *ADE2* via EPIC system. Schematic pJMR19 representing nourseothricin resistance transformation selection marker, Cas9, guide sequence, and autonomous replicating sequence-7 integration into pJMR19. Cas9 mediated disruption of *ADE2* leading to formation of red-pink colonies in a Clade-I *C. auris* isolate, SKU067, on SD agar supplemented with nourseothricin 200mg/L.

To assess incorporation of the *ADE2*^dis^ repair template, 50 red-pink colonies generated in RVA001 and SKU067 were selected for sequencing of the *ADE2* locus (25 per isolate background and representing 3 independent transformations) **(Figure 2A)**. Of the 50 red-pink transformants sequenced, 45 were found to have the targeted manipulation (21 in RVA001 and 24 in SKU067) **(Figure 2B)**. The remaining 5 transformants harbored one of 3 alternative disruptions (**Figure 2B**).

**Figure 2.**
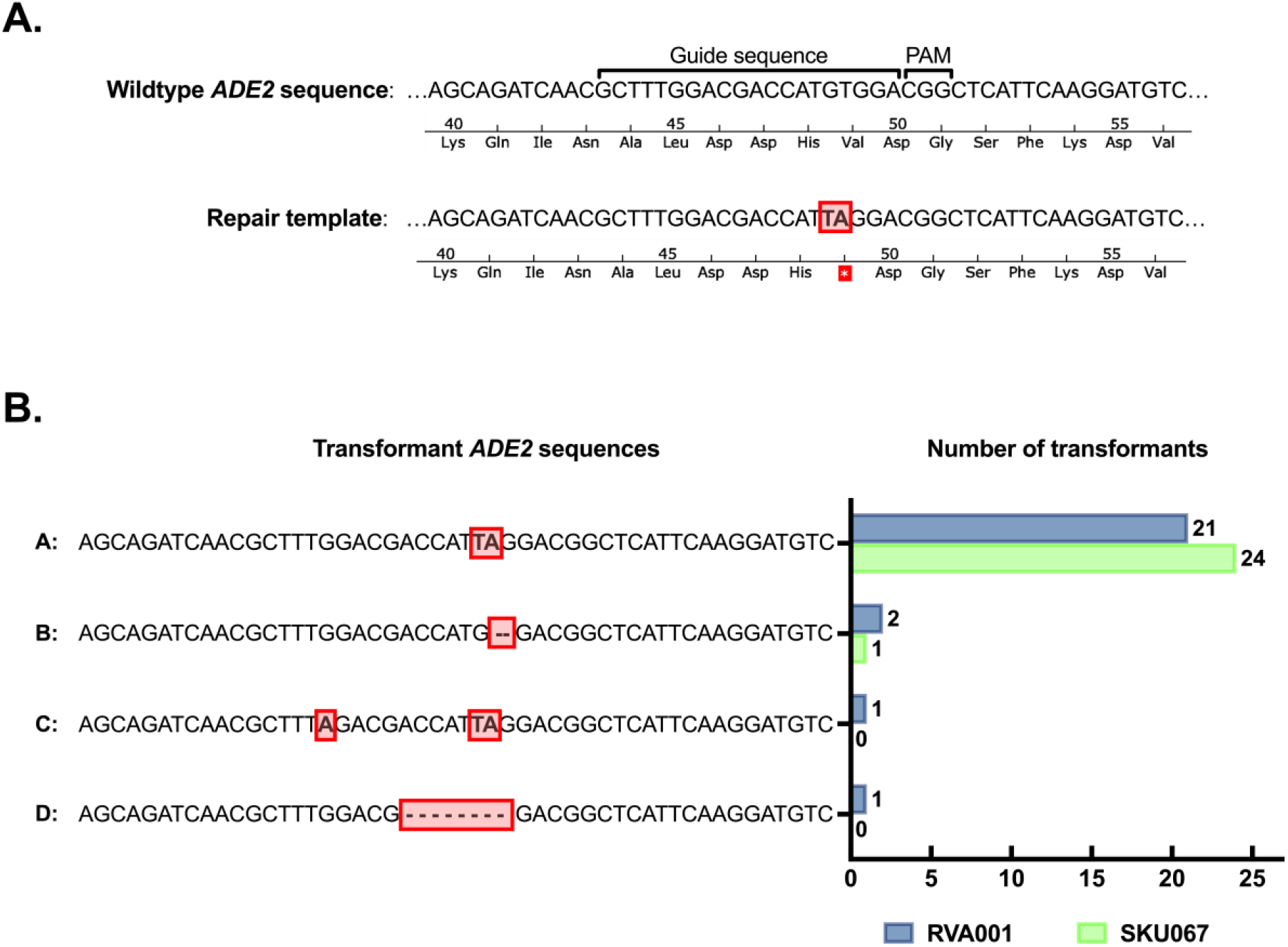
Sequence confirmation of *ADE2* editing. A) Wildtype *ADE2* sequence along with repair template showing intended manipulation highlighted in red to create *ADE2*^dis^ strains. B) Identification and distribution of four variant sequences of *ADE2*^dis^ strains sequenced from SKU067 and RVA001 *ADE2*^dis^ transformations. Variants identified via sequencing by the Center for Advanced Genome Engineering (SJCRH). Plotted values represent distribution for each variant among 50 screened red-pink colonies (25 per isolate background) from 3 independent transformations.

### The *C. auris* EPIC genetic manipulation system is capable of single nucleotide editing

The *C. auris* EPIC system was further characterized by evaluating the direct impact of a single nucleotide *FKS1* mutation frequently associated with clinical echinocandin resistance in the echinocandin-susceptible clinical isolate RVA001 [21]. Transformations were carried out using an *FKS1* hot spot 1 (HS1)-guided EPIC vector (pJMR19-*FKS1*:HS1a) alongside a PCR-generated repair template containing either the *FKS1* sequence encoding an S639F substitution (*FKS1*^S639F^) or the manipulation control sequence (*FKS1*^WTs^; encoding a synonymous nucleotide substitution at codon S639).

Following initial *FKS1* sequencing confirmation, two independent transformants for both RVA001-*FKS1*^WTs^ and RVA001-*FKS1*^S639F^ were passaged in YPD media without nourseothricin to promote loss of the transiently maintained EPIC system vector. gDNA from the 4 transformants and the parental clinical isolate (RVA001) was then extracted for WGS confirmation. Incorporation of the intended single nucleotide substitutions in *FKS1* was found in all 4 transformants. Three of the four transformants were found to have no other nucleotide variations in the genome relative to the parental isolate RVA001, while one strain, RVA001-*FKS1*^WTs^-A, was found to have one additional single nucleotide variation in a non-coding region **(Supplemental Table 2)**.

### The S639F substitution in *C. auris FKS1* confers resistance to clinically available echinocandins

The MIC for all four clinically available echinocandins (anidulafungin, caspofungin, micafungin, and rezafungin) were then assessed for RVA001 and each of the derivative *FKS1* strains. Strains harboring the S639F mutation in *FKS1* were observed to exhibit elevations in MIC for all clinically available echinocandins relative to the parental RVA001 **(Figure 3)**. Anidulafungin MIC increased 64-fold, caspofungin MIC increased 8-fold, micafungin MIC increased 256-fold, and rezafungin MIC increased 16-fold. Conversely, no change in echinocandin MIC was found between RVA001 and the *FKS1*^WTs^ control strains (**Figure 3**).

**Figure 3.**
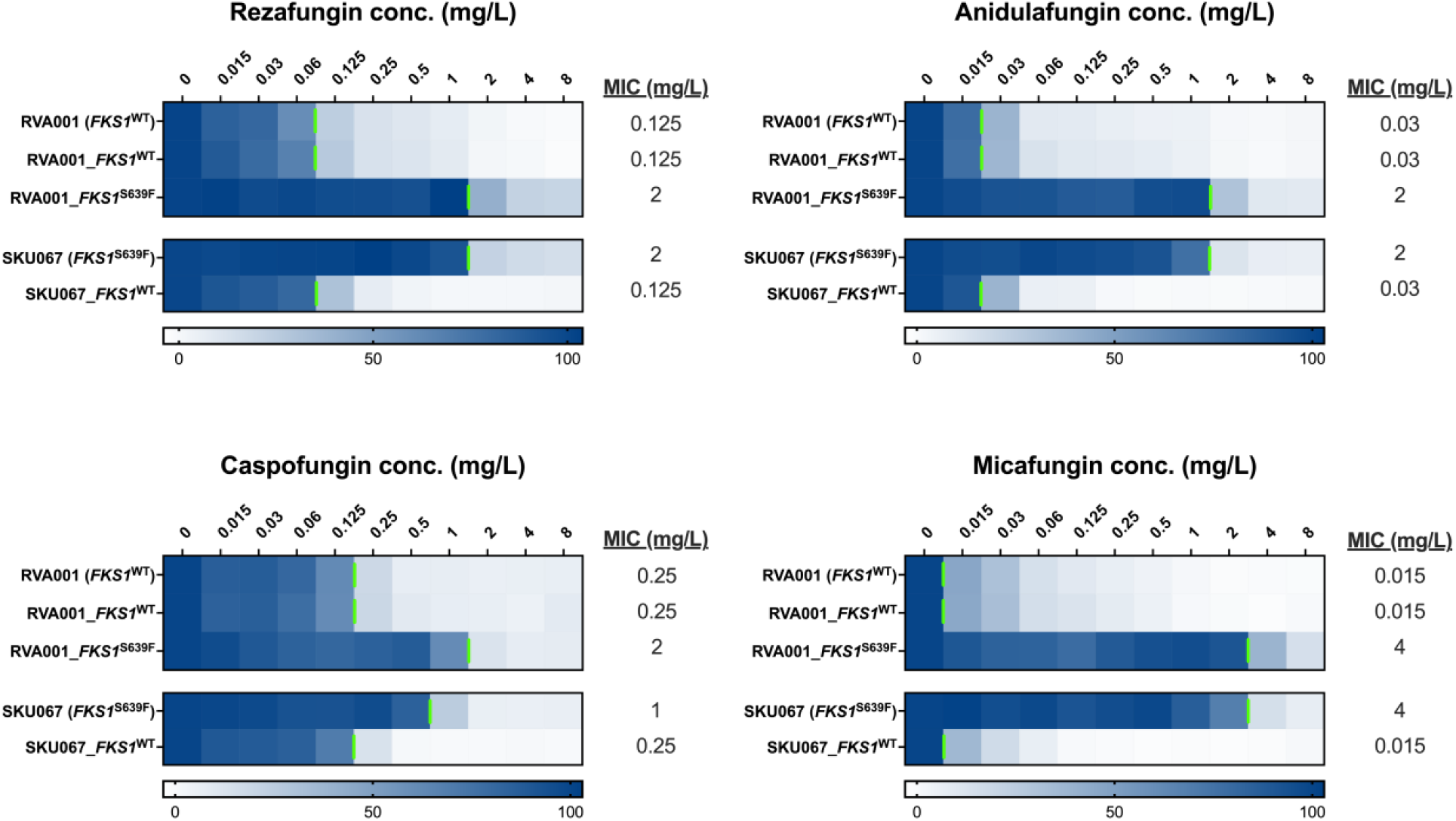
Effects of single *FKS1* mutation on echinocandin susceptibility. Micafungin, anidulafungin, caspofungin, and rezafungin MICs were determined as outlined by the Clinical Laboratory Standards Institute (CLSI) guidelines using broth microdilution. Vertical green lines denote the threshold for 50% growth inhibition compared the growth control well.

To further confirm the impact of the S639F mutation in *C. auris FKS1*, we then used the EPIC system to introduce a sequence encoding wild-type (matching B8441 reference) Fks1 into the echinocandin-resistant clinical isolate SKU067, correcting the S639F substitution. Two independent strains were generated and MIC for all 4 clinically available echinocandins were found to have decreased below current clinical breakpoints (**Figure 3**), definitively demonstrating that the *FKS1*^S639F^ mutation alone is sufficient to confer clinical echinocandin resistance in two independent clinical isolate backgrounds representing both Clade-I and Clade-III.

### The S639F substitution in *C. auris FKS1* confers an increase in the relative abundance of beta-glucans and chitin

Mutations in *FKS1* have previously been associated with altered relative abundance of β-glucan and chitin in *Candida albicans* [22]. We next sought to evaluate the relative abundance of these fungal cell wall constituents in our isogenic sets of *C. auris FKS1* mutation strains by staining with AB and CFW as previously described. Relative abundance of both β-glucan and chitin were observed to increase upon the introduction of the S639F-encoding mutation into both RVA001. Conversely, the abundance of these cell wall constituents also decreased upon correction of the S639F-encoding mutation in SKU067 (**Figure 4**).

**Figure 4.**
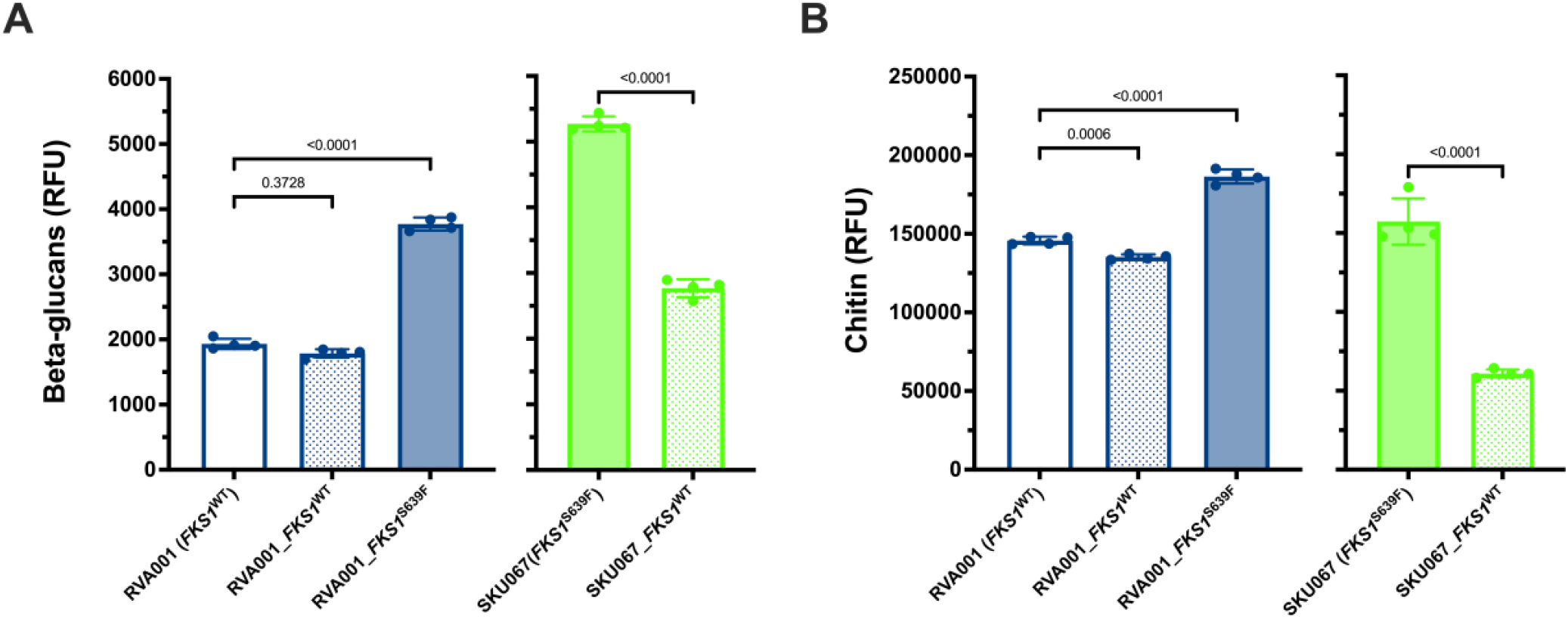
Relative quantification of cell wall components following *FKS1* manipulation via EPIC system. A) Assessment of β-glucan abundance measured by fluorescence intensity of Aniline Blue staining for excitation 395nm and emission 495nm. B) Assessment of chitin abundance measured by CFW fluorescence intensity with excitation 380nm and emission 475nm. P-values derived from ordinary one-way ANOVA statistical analysis.

## DISCUSSION

*C. auris* is a healthcare-associated pathogen of increasing clinical concern and represents a substantial threat to global public health. While the growing body of research has revealed much about *C. auris*, a lack of available molecular tools impedes progress and many important questions pertaining to this organism remain unanswered. The *C. auris-*optimized EPIC system described here represents an important new genetic tool for the study of *C. auris*. In addition to the precise single-nucleotide editing of *ADE2* and *FKS1*, the EPIC system can also be used to introduce larger gene cassettes, such as those expressing fluorescent proteins, as demonstrated by the generation of a dTOM-fluorescent strain in the derived RVA001-*FKS1*^S639F^ strain **(Supplemental Figure 2)**. Further demonstrating the utility and recyclability of the EPIC system. Using this novel tool, we show the impact of one of the most commonly observed *C. auris*

*FKS1* mutations associated with clinical echinocandin resistance, encoding S639F. Demonstrated by manipulations both introducing and correcting this mutation in two independent clinical isolate backgrounds representing two different genetic clades of *C. auris*, we have delineated the direct contribution of the mutation encoding S639F to clinical echinocandin resistance. The S639F mutation alone is sufficient to confer increased resistance to all 4 clinically available echinocandins, increasing echinocandin MIC by as much as 256-fold and elevating rezafungin and micafungin MIC above available clinical breakpoints in both clinical isolate backgrounds tested [3, 23]. Further, using our isogenic sets of *FKS1* mutant strains, we have shown this mutation also alters the relative abundance of both chitin and β-glucans, suggesting the impact of these mutations may extend beyond impeding echinocandin interaction with Fks1.

Taken together, the findings of this study definitively demonstrate the S639F-encoding *FKS1* mutation confers dramatically increased resistance to clinically available echinocandins, including the recently approved agent rezafungin. Further, we show the potential utility of a novel *C. auris*-optimized EPIC gene-editing system, a recyclable tool capable of single nucleotide editing which we hope will enable future research seeking to characterize and overcome *C. auris* and the infections caused by this emerging pathogen of global concern.

## NOTES

### Transparency declaration

No reported conflicts. Authors have submitted the ICMJE Form for Disclosure of Potential Conflicts of Interest. Conflicts that the editors consider relevant to the content of the manuscript have been disclosed. This work was supported through the St. Jude Children’s Research Hospital Children’s Infection Defense Center grant and the Society of Infectious Diseases Pharmacists Young Investigator Research Award granted to J.M.R.. The funders had no role in study design, data collection and interpretation, or the decision to submit the work for publication.

## Acknowledgments

This research was supported in part by ALSAC and the National Cancer Institute grant P30 CA021765. Preliminary results of these studies were presented on the 23^rd^ of April, 2022 at the “New mechanisms of antifungal resistance in emerging yeast pathogens” oral session as part of the 32nd European Congress of Clinical Microbiology & Infectious Diseases with the support of the 32nd ECCMID 2022 Travel Grant. The authors would like to thank Lisa Lombardi and Geraldine Butler for kindly providing the pCP-tRNA vector, and Lisa Lombardi, Geraldine Butler, and Judith Berman for helpful discussions relating to this manuscript.

