## Supplementary Data for "A *Candidozyma* (*Candida*) *auris*-optimized Episomal Plasmid Induced Cas9-editing system reveals the direct impact of the S639F encoding *FKS1* mutation"

### Supplemental Table 1: Isolates, Strains, and EPIC components

| Isolates and Strains | Description |
|---|---|
| AR0931 | Clade IV isolate |
| AR1097 | Clade V isolate |
| AR1100 | Clade II isolate |
| SKU067 | Echinocandin resistant Clade I isolate |
| RVA001 | Echinocandin susceptible Clade III isolate |
| RVA001-FKS1$^{WTs}$-A | RVA001 derived manipulation control strain containing synonymous C1917A mutation |
| RVA001-FKS1$^{WTs}$-B | RVA001 derived manipulation control strain containing synonymous C1917A mutation |
| RVA001-FKS1$^{S639F}$-A | RVA001 derived strain containing non-synonymous C1916T mutation |
| RVA001-FKS1$^{S639F}$-B | RVA001 derived strain containing non-synonymous C1916T mutation |
| Kw2999-FKS1$^{WT}$-A | SKU067 derived strain containing non-synonymous T1916C mutation and synonymous TTGTCC 1939..1944 CTTCTG substitution |
| Kw2999-FKS1$^{WT}$-B | SKU067 derived strain containing non-synonymous T1916C mutation and synonymous TTGTCC 1939..1944 CTTCTG substitution |
| RVA001-FKS1$^{S639F}$-A-dTom | RVA001 derived strain containing non-synonymous C1916T mutation and the fluorescent dTOM expression cassette inserted at base 107955 in chromosome 6 |

| Plasmid | Description |
|---|---|
| pJMR17v3 | Rybak JM, Barker KS, Muñoz JF, et al. In vivo emergence of high-level resistance during treatment reveals the first identified mechanism of amphotericin B resistance in *Candida auris*. Clin Microbiol Infect. 2022;28(6):838-843. doi:10.1016/j.cmi.2021.11.024 |
| pJMR19 | *C. auris*-optimized **E**pisomal **P**lasmid **I**nduced **C**as9 (EPIC) system vector |
| pKE4-dTOM | Butts A, DeJarnette C, Peters TL, et al. Target Abundance-Based Fitness Screening (TAFiS) Facilitates Rapid Identification of Target-Specific and Physiologically Active Chemical Probes. mSphere. 2017;2(5):e00379-17. Published 2017 Oct 4. doi:10.1128/mSphere.00379-17 |

| Primer | Sequence | Description |
|---|---|---|
| **pJMR19 build** | | |
| CAU-ACT1-F | tggtACGCGTGAATTCTTCGATAAAAGTAGGAAGAAAGAG | Amplification of ACT1 promotor for *C. auris* optimization of pJMR17v3 |
| CAU-ACT1-R | TGGTGGTACCCAATGAATGTACTTAAGCTCGC | Amplification of ACT1 promotor for *C. auris* optimization of pJMR17v3 |
| pJMR17v3 AMP-F | tggtACGCGTatggacggtgaagaagttgctgctttagttatcg | Amplification of vector backbone from pJMR17v3 |
| pJMR17v3 AMP-R | TGGTGGTACCgttttcccagtcacgacgtt | Amplification of vector backbone from pJMR17v3 |
| ***ADE2* Modification** | | |

| | | |
|---|---|---|
| *ADE2*-gBlock for Repair Template PCR | CCCACTAATAGCTTTTCGCAGCCATTAATTGAAGCTACTATGAGCATCAACATACTCAGCAAGTACACAACTTG<br>GAAAAAAATAAGAAATGAGCTTAATTAAAGTTGGCCCTCCCAAATAATCAAAATCTCGCCAGAATATTCATAATT<br>CCTTCTTCTACAGTCATTATCTTGCACTCGCCAACATCAGCGAGATGCCCCATCGATGTCACTGCACCTCTGAC<br>TCTGTTCGCATCTCACATCACCAACTACAACTACCATCTTCATTATCATGGACGGAAAAACAATTGGAATTCTTG<br>GAGGCGGCCAGTTGGGGCGGATGATTGTTGAGGCTGCACACAGACTCAACATTAAAACTGTGGTGCTTGACG<br>CTGCTCATTCTCCAGCCAAGCAGATCAACGCTTTGGACGACCATGTGGACGGCTCATTCAAGGATGTCGATGC<br>TATCACCAGGTTGGCCAAAAAATGTGATTTGTTGACGATAGAGATTGAGCACGTTGACGTGGAGGCGTTGAAG<br>ACGGTCGAAAAGACCTACAATGTGCCCATCTACCCGTTCCCAGAGACAATCAGGCTCATCCAGGATAAGTACC<br>TTCAAAAGACCCATTTGATGGAGCACGGTGTTGATGTGGTTGAGTCGATTGCCGTGAAAGAAAACACCAAGGA<br>GGGTTTGCAAGAAATTGGCGCTAAGCTTGGCTTTCCCTTTATGTTGAAGTCGAGAACGATGGCCTACGACGGT<br>AGAGGAAACTTCGTTGTGAAGAATGAAGAGGCCATTCCTGAGGCGCTAGAATTTTTGTCCAACAGACCTCTTTA<br>CGCGGAAAAATGGTGCCCTTTCACCAAAGAATTGGCAGTGATGGTGGTTAGACTGCTTGAAGGCGAGGTGTTT<br>GCGTATCCTACTGTTGAAACCCACCACAAGGATAACATATGCCATTTGGTTTACGCTCCTGCCAGAGTCCTGG<br>ACACTTTGCAGAAGAAAGCCTCAATATTGGCGCAAAACGCCGTCAAGTCCTTCCC | |
| *ADE2*-top | ccaGCTTTGGACGACCATGTGGA | Guide sequence inserted into pJMR19 editing of ADE2 |
| *ADE2*-bot | aacTCCACATGGTCGTCCAAAGC | Guide sequence inserted into pJMR19 editing of *ADE2* |
| *ADE2*-RT-F | CCCACTAATAGCTTTTCGCAGCC | Amplification of *ADE2* gblock |
| *ADE2*-RT-R | GGGAAGGACTTGACGGCG | Amplification of *ADE2* gblock |
| *ADE2*-Screen-F | CCCATCGATGTCACTGCACC | PCR primer for *ADE2*$^{dis}$ MiSeq |
| *ADE2*-Screen-R | CCTGGATGAGCCTGATTGTCTC | PCR primer for *ADE2*$^{dis}$ MiSeq |

**<u>*FKS1* Modification</u>**

| | | |
|---|---|---|
| *FKS1*-HS1a-top | ccaGTTTCTAATAGGATCTCTCA | Guide sequence inserted into pJMR19 for editing of *FKS1* in RVA001 |
| *FKS1*-HS1a-bot | aacTGAGAGATCCTATTAGAAAC | Guide sequence inserted into pJMR19 for editing of *FKS1* in RVA001 |
| *FKS1*-HS1b-top | ccaATGGTGGACAAGTTTCTAAT | Guide sequence inserted into pJMR19 for editing of *FKS1* in SKU067 |
| *FKS1*-HS1b-bot | aacATTAGAAACTTGTCCACCAT | Guide sequence inserted into pJMR19 for editing of *FKS1* in SKU067 |
| *FKS1*-HS1a-RT-WTs-F | CCTGGTGTTTACACAAGGTATCACCA<br>AACCATTGTTCACCGTTGCATCTCAT<br>CGTCATGGTGGACAAGTTTCTAATAG<br>GA | Amplification of Repair template: *FKS1* Silent Mutation in RVA001 |
| *FKS1*-HS1a-RT-WTs-R | ACTGTGTTTGCTGCTAAGTTGGCCG<br>AATCTTACTTCTTCTTGACTTTGTCaT<br>TGAGAGATCCTATTAGAAACTTGTCC<br>AC | Amplification of Repair template: *FKS1* Silent Mutation in RVA001 |
| *FKS1*-HS1a-RT-S639F_F | GGTGGCCAAAGACTGAGCAAAG | Amplification of *FKS1*$^{S639F}$ repair template sequence from SKU067 gDNA |
| *FKS1*-HS1a-RT-S639F-R | CCGCATCTTCGCAATGGTC | Amplification of *FKS1* $^{S639F}$ repair template sequence from SKU067 gDNA |
| *FKS1*-HS1b-RT-WT-F | CCTGGTGTTTACACAAGGTATCACCA<br>AACCATTGTTCACCGTTGCATCTCAT<br>CGTCATGGTcagaagGTTTCTAATAG | Amplification of Repair template: *FKS1*$^{WT}$ Silent Substitutions in SKU067 |
| *FKS1*-HS1b-RT-WT-R | TGCTGCTAAGTTGGCCGAATCTTACT<br>TCTTCTTGACTTTGTCCTTGAGAGAT<br>CCTATTAGAAACcttCTGACCATGACG<br>A | Amplification of Repair template: *FKS1*$^{WT}$ Silent Substitutions in SKU067 |
| *FKS1*-ORF-Screen-F | CGCTACCTTGCTTTTCTTCGC | PCR amplification primer for *FKS1* sequencing |
| *FKS1*-ORF-Screen-R | CTCCATCTCTGTGGTGGCC | PCR amplification primer for *FKS1* sequencing |

**<u>dTOM Insertion</u>**

| | | |
|---|---|---|
| dTOM landing-pad-top | ccaCTTTCAAACGAAGGCTGCGG | Guide sequence inserted into pJMR19 for insertion of dTOM expression cassette in chromosome 6 |
| dTOM landing-pad-bot | aacCCGCAGCCTTCGTTTGAAAG | Guide sequence inserted into pJMR19 for insertion of dTOM expression cassette in chromosome 6 |
| dTOM landing-pad-RT_F | CTCTGTGTCGCATGGAGACAATGGG CCCCATGAAGAGGACACAGGGAGGT CTGAAAAGAAAAAAATTAGTGAGGGT ACCTGCAAATCTGTTTG | Amplification of dTOM from pKE4 |
| dTOM landing-pad-RT_R | CTGGCAATAAGCAACCTCCCCAGGA AAACCCATGACCTTTTCACGGACCTT CAACCCGCCGTTCTCACCACCTTCT AGATACCTAGGTGAGCTC | Amplification of dTOM from pKE4 |
| Landing-pad-Screen-F | GCGGAAGCAAGCTCAGGG | PCR primer for Landing Pad dTOM screening |
| Landing-pad-Screen-R | CGAGGAATCCCACGGGAGG | PCR primer for Landing Pad dTOM screening |

**Supplemental Table 2. Variant summary via whole genome sequencing analysis**

| Sample | Variants per region | | Coding region variant type |
|---|---|---|---|
| | Coding region | Non-Coding Region | |
| RVA001 | 0 | 0 | N/A |
| RVA001-*FKS1*$^{S639F}$-F1A | 1 | 1 | synonymous_variant: *FKS1* |
| RVA001-*FKS1*$^{S639F}$-O1A | 1 | 0 | synonymous_variant: *FKS1* |
| RVA001-*FKS1*$^{WT}$-216A | 1 | 0 | missense_variant: *FKS1* |
| RVA001-*FKS1*$^{WT}$-L1A | 1 | 0 | missense_variant: *FKS1* |

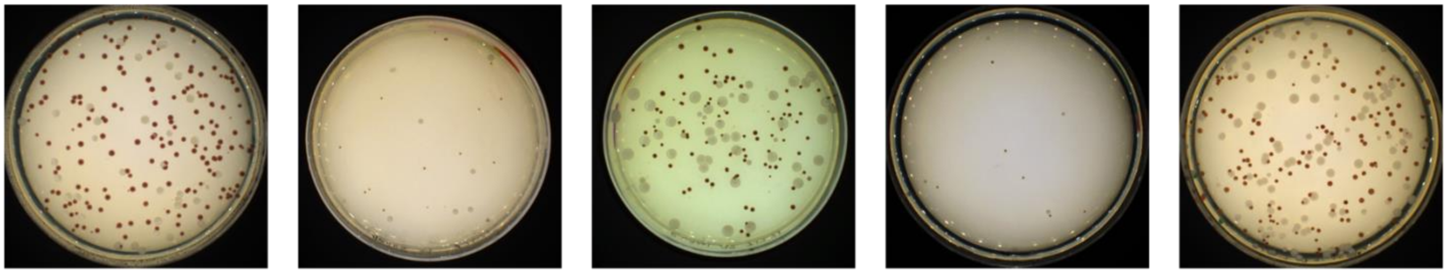

**SKU067 (Clade I)** | **AR1100 (Clade II)** | **RVA001 (Clade III)** | **AR0931 (Clade IV)** | **AR1097 (Clade V)**

**SUPPLEMENTAL FIGURE 1. EPIC mediated *ADE2* editing in 5 subclades of *C. auris.***
Presence of red pigmented colonies indicates successful editing of the *ADE2* locus. All transformants plated onto SD media containing 200mg/L nourseothricin for EPIC vector presence selection.

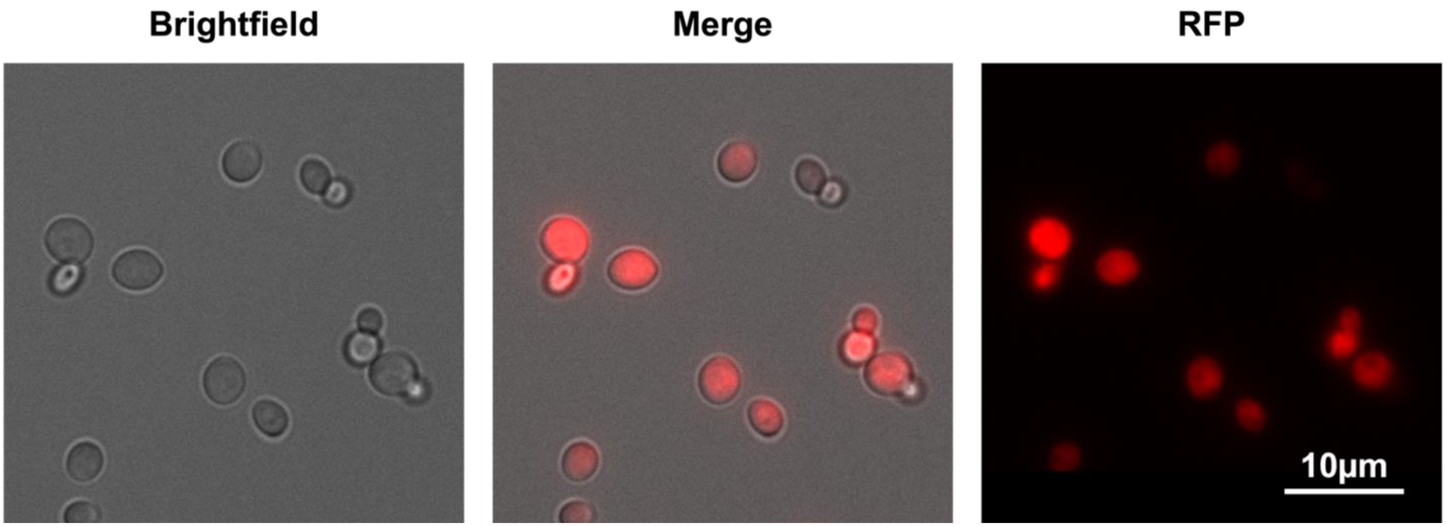


**Supplemental Figure 2. EPIC mediated insertion of a dTOM cassette in the derived RVA001-FKS1$^{S639F}$ strain.** RVA001-*FKS1*$^{S639F-}$dTom was generated using the EPIC vector carrying a guide sequence (Supplemental Table 1) targeting the midpoint of a 10kb region of Chromosome 6 with no predicted open reading frames and a dTom expression cassette amplified by PCR from the PKE4-dTOM vector using primers (Supplemental Table 1) which introduced ~70mer of homology matching the sequences flanking the intended Cas9 cut site. Samples grown overnight at 35°C, 220RPM in YPD liquid media. Cells were pelleted at 4000 RPM and washed 2x with sterile PBS before brightfield and RFP2.0 imaging at 60x magnification on an EVOS M5000 (Invitrogen). Individual image file alignment and color overlay was performed using ImageJ.
